# Development of an MRI-compatible robotic perturbation system for studying the task-dependent contribution of the brainstem to long-latency responses

**DOI:** 10.1101/2024.03.01.583025

**Authors:** Rebecca C. Nikonowicz, Fabrizio Sergi

## Abstract

Methodological constraints have hindered direct *in vivo* measurement of reticulospinal tract (RST) function. The RST is thought to contribute to the increase in the amplitude of a long latency response (LLR), a stereotypical response evoked in stretched muscles, that arises when participants are asked to “resist” a perturbation. Thus, functional magnetic resonance imaging (fMRI) during robot-evoked LLRs under different task goals may be a method to measure motor-related RST function. We have developed the Dual Motor StretchWrist (DMSW), a new MR-compatible robotic perturbation system, and validated its functionality via experiments that used surface electromyography (sEMG) and fMRI. A first study was conducted outside the MRI scanner on six participants using sEMG to measure wrist flexor muscle activity associated with LLRs under different task instructions. Participants were given a Yield or Resist instruction before each trial and performance feedback based on the measured resistive torque was provided after every “Resist” trial to standardize LLR amplitude (LLRa). In a second study, ten participants completed two sessions of blocked perturbations under 1) Yield, 2) Resist, and 3) Yield Slow task conditions (control) during whole-brain fMRI.

Statistical analysis of sEMG data shows significantly greater LLRa in Resist relative to Yield. Analysis of functional images shows increased activation primarily in the bilateral medulla and midbrain, and contralateral pons and primary motor cortex in the Resist condition. The results validate the capability of the DMSW to elicit LLRs of wrist muscles with different amplitudes as a function of task instruction, and its capability of simultaneous operation during fMRI.

## I. INTRODUCTION

The reticulospinal tract (RST), while secondary to the corticospinal tract (CST) in healthy participants, may assume considerable importance for recovery from corticospinal lesions [1], [2]. The RST originates in the brainstem and is primarily thought to control gross movements such as locomotion and posture, but we have recently discovered that it contributes to reaching and grasping movements [3]. There is also evidence that the RST can strengthen its outputs in the event of a corticospinal lesion, though it appears there are limitations in the degree of recovery, specifically the level of dexterity, that can be achieved with RST contribution [4], [5], [6]. Methodological constraints have limited our capability to directly measure *in vivo* function in the brainstem nuclei originating from the RST, resulting in an incomplete understanding of the role of RST in neuromotor impairment and recovery. Advances in neuroimaging provide an opportunity to measure task-related neural activity *in vivo* in the brainstem, a structure that is small in size and located deep in the cranium [7], [8].

Recent studies have shown that the Reticular Formation (RF), a set of brainstem nuclei that originate the RST, may be actively involved in the generation of long latency responses (LLRs), a stereotypical response evoked in muscles stretched by a perturbation [9]. LLRs, occurring 50-100 ms post perturbation onset, offer a reliable and less confounded measure of RST function. Their stereotypical nature ensures greater consistency across trials and participants compared to voluntary tasks, enhancing the specificity of RST-related activity. Perturbation-induced LLRs could serve as a dependable stimulus for directly evaluating motor activity in the RF during rapid joint stretch using neuroimaging [10].

Our lab has recently developed StretchfMRI, a novel technique to study the brainstem correlates of LLRs *in vivo* in humans [10] StretchfMRI combines robotic perturbations with surface electromyography (EMG) and functional Magnetic Resonance Imaging (fMRI) to provide simultaneous recording of neural and muscular activity associated with LLRs. Preliminary studies have validated the technique and shown that neural activity associated with LLRs in the flexor carpi radialis and extensor carpi ulnaris can be reliably measured in the RF [10]. While StretchfMRI has shown promise, it has insofar been limited in its functional specificity, making it challenging to discern whether observed neural activation is due to background activity or a response to stretch-induced reflex responses. To advance our understanding of RST function, it is crucial to decouple background activation from activation solely attributable to LLRs and to decouple RF activation due to CST engagement versus RST engagement. Because the RST is thought to contribute to the increase in LLR amplitude (LLRa) associated when participants are asked to “resist” a perturbation [11], fMRI during LLRs under different task goals may be a highly specific method to decouple the contribution of the CST and RST to LLRs. We have extended the capabilities of StretchfMRI to measure LLR-related fMRI signal under different task instructions, designed to differentially engage the CST and RST, by building a new MR-compatible robotic perturbator, the Dual Motor StretchWrist (DMSW). In this paper, we present the design of the DMSW and we describe a new study protocol used to condition long-latency responses under different task instructions, that can be executed within the performance constraints imposed by the requirement of MRI-compatibility. Moreover, we demonstrate that the developed system is able to elicit LLR of wrist muscles with different amplitude as a function of task instructions, and of operating simultaneosuly during fMRI.

## II. METHODS

### A. Dual Motor StretchWrist Development

Our lab has previously developed the MR-StretchWrist, an MR-compatible 1-degree-of-freedom robot, capable of delivering perturbations to the wrist joint of participants asked to “yield” to the perturbation [10]. However, the MR-StretchWrist cannot operate in conditions during which participants “resist” a perturbation due to performance limitations of the piezoelectric ultrasonic motor used (EN60 motor, Shinsei Motor Inc., Japan) and of the capstan transmission. In fact, the reaction torque produced by participants exceeds the 3 Nm peak torque that can be delivered by the MR-StretchWrist, thus causing the motor to lock and the driver to go into protection.

To enable our task-dependent paradigm with a “resist” condition, we have developed a new MRI-compatible 1degree-of-freedom wrist robot, the Dual Motor StretchWrist (DMSW), capable of imposing controlled perturbations to the hand about the wrist flexion/extension axis in the range of *θ*_*F E*_ = [-45, 45]° with up to 6 Nm of peak torque (Fig. 1). It is actuated by two ultrasonic piezoelectric motors connected in parallel (EN60 motor, Shinsei Motor Inc., Japan) via a capstan transmission with a 3:1 gear ratio to transfer motion from the motors to the end effector. The capstan drive consists of 3 pulleys with different diameters connected with a smooth cable wrapped multiple times to ensure no slippage, high friction contact.

**Fig. 1.**
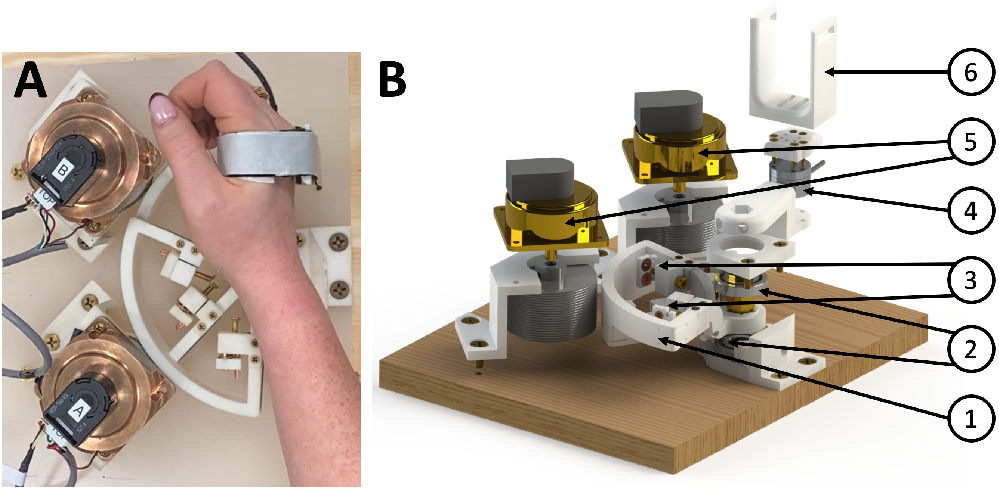
**(A)** Dual Motor StretchWrist top view with a participant’s hand. **(B)** Exploded view of the Dual Motor StretchWrist. (1) Output capstan arc, (2) structural ceramic bearings, (3) tensioning mechanisms, (4) MR-compatible force/torque sensor, (5) ultrasonic motors, and (6) hand support.

To ensure MR-compatibility of the robot, all structural components are manufactured using ABS-based 3D-printed plastic (RS-F2-FPWH-04, Formlabs, Inc., MA, USA) and connected using brass screws. The capstan transmission cable is a microfiber braided wire (SpiderWire Stealth 0.4 mm diameter braided fishing line, 80 lb test). The output shaft is supported by ceramic radial bearings (Boca Bearings, Boynton Beach, FL, USA) and bronze sleeve bearings are used to support the tensioning mechanisms. Electromagnetic interference is reduced via a hexapolar twisted-pair shielded cable for the motor encoder line and a shielded cable for the motor power line. These lines are filtered when passing through the scanner patch panel using 5.6 pF and 1.3 pF capacitive filters, respectively. To measure torque produced by the participant’s wrist, we have included a six-axis MR-compatible force/torque sensor (Mini27Ti, ATI Industrial Automation, Apex, NC).

A controller ensures proper parallel function of the motors, including matching motor displacements and velocities. Due to properties inherent to the individual motors and drivers, the same voltage delivered to two motors will result in slightly different velocity profiles. Thus, the P controller used for the StretchWrist was augmented for the DMSW such that the desired velocity command for each motor is updated separately, with the feed-forward command

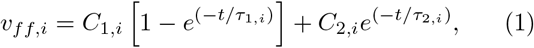

with gains and time constants tuned to the corresponding motor/driver (Fig. 2A). These variables were tuned via an iterative method until both motors were moving with *<*1° difference in angular displacement at all times during a perturbation (Fig. 2B).

**Fig. 2.**
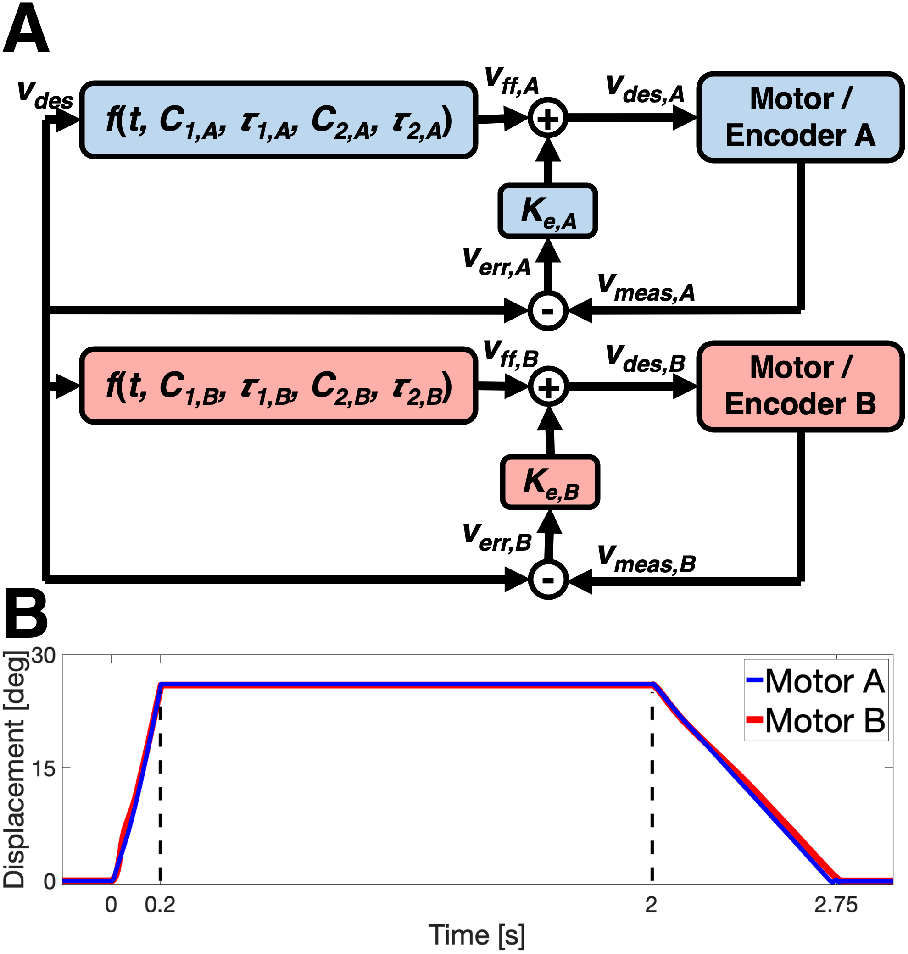
**(A)** Block diagram of the DMSW’s P controller. **(B)** Motor displacements showing *<*1° difference in angular displacement at all times during a perturbation.

### B. Protocol Validation

To confirm the DMSW’s ability to reliably elicit LLRs during both “Yield” and “Resist” conditions, a preliminary study was conducted outside the MRI scanner on 6 healthy, right-handed participants using surface EMG to measure muscle activity. The experiment consisted of 4 sessions: Familiarize 1, Familiarize 2, Task 1, and Task 2, with rests between each session as needed. During all sessions, participants completed five 1 s isometric contractions corresponding to 0.2 Nm of torque in extension followed by a series of ramp-and-hold 200 ms perturbations, in extension. During perturbations, participants were provided with a visual target corresponding to 0.2 Nm of torque in flexion as a background torque, followed by either a 150 deg/s or 35 deg/s perturbation. Participants were given a “Yield” or “Resist” instruction prior to each trial.

During Familiarize sessions, participants completed the 4 block conditions described in Table I. To standardize the resistance that participants needed to apply during a perturbation, visual performance feedback was provided based on average torque measured during a time interval *δ*T=[75-125] ms from perturbation onset, assumed to result from muscle activation applied in the long-latency window. For Familiarize sessions, performance feedback was provided only after Resist trials to match 133% of the average torque measured for each perturbation during the previous Yield block in the same interval. Each block occurred once and consisted of 20 trials.

**TABLE I.** FAMILIARIZE SESSION BLOCKS.

| Block ID | Bkgd. Torque | Inst. | Vel. | Perf. Feedback |
| --- | --- | --- | --- | --- |
| Relax | No | Yield | 150 deg/s | No |
| Yield | Yes | Yield | 150 deg/s | No |
| Resist | Yes | Resist | 150 deg/s | Yes |
| Yield Slow | Yes | Yield | 35 deg/s | No |

During Task sessions, participants completed the 3 block conditions described in Table II. For Task sessions, performance feedback was provided after every trial based on the average torque measured during Familiarize 2. Task Yield feedback goal was 100±15% of the Familiarize 2 Yield average, Task Resist was 133±15% of the Familiarize 2 Yield average, and Task Yield Slow was 100±15% of the Familiarize 2 Yield Slow average. Each block occurred 6 times and consisted of 7 trials for a total of 126 perturbations. Block order was randomized but held consistent between participants.

**TABLE II.** TASK SESSION BLOCKS.

| Block ID | Bkgd. Torque | Inst. | Vel. | Perf. Feedback |
| --- | --- | --- | --- | --- |
| Yield | Yes | Yield | 150 deg/s | Yes |
| Resist | Yes | Resist | 150 deg/s | Yes |
| Yield Slow | Yes | Yield | 35 deg/s | Yes |

Participants were lying in a supine position in a mock MRI scanner with the DMSW secured on a wooden box above their torso (Fig. 3). Electrodes (Delsys Trigno Avanti, Natrik, MA) were placed on the flexor carpi radialis and the extensor carpi ulnaris to measure muscle activity.

**Fig. 3.**
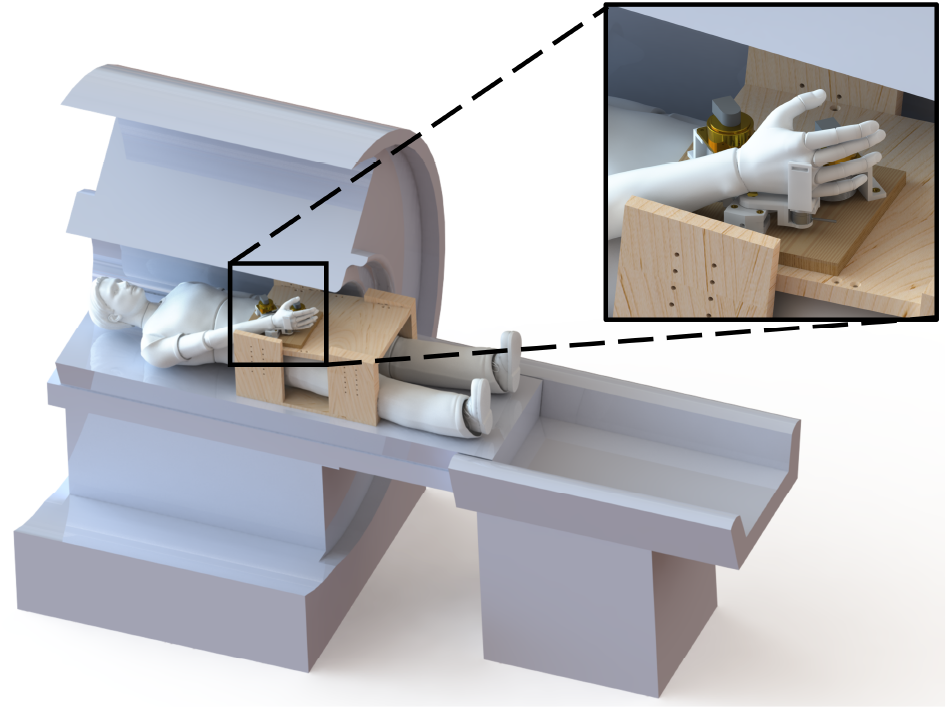
Experimental setup with the participant in a supine position inside the MRI scanner with the DMSW resting on top of a wooden box.

### C. fMRI Pilot Experiment

12 healthy individuals (7 males, 5 females; age range 22-31 years) were recruited for a pilot study. All participants self-reported as right-handed, free from neurological disorders, orthopedic or pain conditions affecting the right arm, and provided written consent prior to data collection. The study was approved by the Investigation Review Board at the University of Delaware under protocol number 1097082-8. Of the 12 participants consented, 2 participants were not able to complete the experiment due to technological difficulties. As a result, fMRI analysis was performed with 10 participants.

Participants were screened for MRI safety and then completed a series of familiarization tasks with the robot to determine their personalized torque target values and ensure they were comfortable with the device and the task instructions. Participants were then positioned in the scanner bore with the robot and completed two Task sessions of blocked perturbations during fMRI sequencing lasting approximately 15 min each. Conditions included were 1) Yield, 2) Resist, and 3) Yield Slow, interleaved with 15 s rest periods.

Experiments were conducted at the University of Delaware Center for Biomedical and Brain Imaging with a Siemens Prisma 3T scanner and a 64 channel head coil. fMRI images were collected using a whole-brain sequence (MultiBand Accelerated EPI with 2×2×2 mm^3^ voxel resolution, TR=1000 ms), together with a T1-weighted structural scan (0.7×.0.7×0.7 mm^3^, TR=2300 ms) used for registration and normalization. Participant pulse and respiration were recorded during the experiment via an MR-compatible pulse oximeter and a respiration belt (Siemens Physiology Monitoring Unit - PMU).

### D. Statistical Analysis of EMG and Torque Data

Raw EMG data were pre-processed using a standard pipeline: band-pass filter to remove motion artifacts and high frequency noise (4th order Butterworth filter), signal rectification, and low-pass filter to extract the envelope of the EMG signal (4th order Butterworth filter).

EMG signal for FCR was normalized by dividing the pre-processed signal by the average pre-processed signal measured in the last 200 ms during the background contraction before perturbation onset across all perturbations. EMG signal for ECU was normalized by dividing the pre-processed signal by the average pre-processed signal measured in the five 1-s long isometric contractions conducted at the beginning of each experiment. LLR amplitude (LLRa) for each trial was defined as the average FCR EMG signal in the [50-100] ms window post perturbation onset. Torque data from the DMSW’s force-torque sensor were processed in Matlab 2023a (Mathworks, Inc., Natick, MA, USA). Torque response amplitude was quantified as the average torque in the time interval *δ*T=[75-125] ms from perturbation onset.

EMG and torque data were analyzed using a linear mixed model with instruction as a main effect and, for group analysis, participant as a random effect. using JMP analysis software (Version Pro 17, SAS Institute Inc., Cary, NC).

### E. Statistical Analysis of Imaging Data

Analysis of all fMRI datasets was performed using SPM12 (Wellcome Department of Cognitive Neurology, London, UK) running on Matlab 2023a. Functional data were preprocessed using a standard pipeline: realignment to the mean image, co-registration to structural MPRAGE, normalization to a standard MNI space template, spatial smoothing, and high-pass filtering (cut off at 128 s). A Gaussian kernel with FWHM=4 mm for brainstem-specific analysis and FWHM=8 mm for whole-brain analysis were used for spatial smoothing.

First-level analyses were performed separately to determine whole-brain and brainstem-specific activation. The same general linear model (GLM) was used for both analyses, however, the model was convolved with a standard hemodynamic response function (HRF) for the whole-brain analysis and a brainstem-specific HRF for assessing activation in the brainstem. The neural response was modeled as

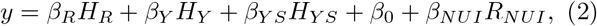

where *H*_*R*_, *H*_*Y*_, and *H*_*Y S*_ are the task regressors (rectangular functions convolved with a HRF) and *R* is a set of 24 nuisance regressors (18 physiological noise regressors and 6 head movement regressors computed using the RETROICOR algorithm via the TAPAS PhysIO Toolbox [12]. GLM estimation was used to identify voxels whose signal was significantly modulated by Yield Slow blocks (*β*_*Y S*_ *>* 0), Yield blocks (*β*_*Y*_ *>* 0), and Resist blocks (*β*_*R*_ *>* 0). Then, participant-specific statistical parametric maps were used as inputs to determine group-level effects in second level analyses.

## III. RESULTS

### A. Protocol Validation

Statistical analysis of sEMG data collected during the protocol validation experiment (Fig. 4) shows significantly greater LLRa in Resist relative to Yield for all individual participants and at a group level. LLRa for Yield Slow was not significantly lower compared to LLRa for Yield at the group level, but it was significantly lower for 4/6 participants (Group Resist, mean ± s.e.m.: 5.7 ± 0.8 n.u., Group Yield: 1.5 ± 0.8 n.u. Group Yield Slow: 1.0 ± 0.8 n.u., Group *p* = 0.0094).

**Fig. 4.**
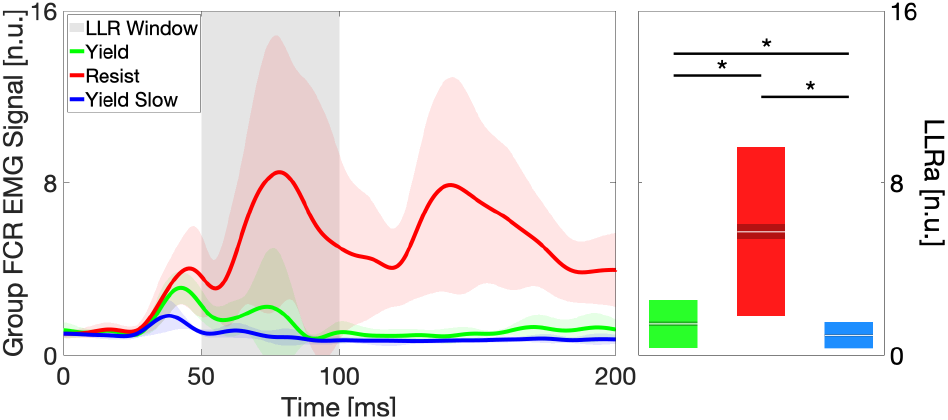
Group-average of the LLRa measured in different instruction conditions in response to a 150 deg/s ramp-and-hold perturbation. The thick line represents the group mean and the shaded area ± one standard deviation (s.d.) of the group mean. The gray shaded area *±* indicates the time interval where an LLR is expected. Box plots indicate the distribution of LLR amplitude, with indications for the mean (white horizontal line), mean *±* one standard error of the mean (dark shaded area), and mean *±* one s.d. (light shaded area).

Statistical analysis of torque data shows significantly greater average torque measured in the [75-125] ms window post-perturbation onset for Resist compared to Yield at a group level and for all participants at an individual level (Fig. 5). Additionally, Yield torque averages were greater than Yield Slow torque averages at a group level and for all participants at an individual level (Group Resist: 0.51 ± 0.03 Nm, Group Yield: 0.40 ± 0.03 Nm, Group Yield Slow: 0.27 ± 0.03 Nm, *p <* 0.001 for all pair-wise comparisons.

**Fig. 5.**
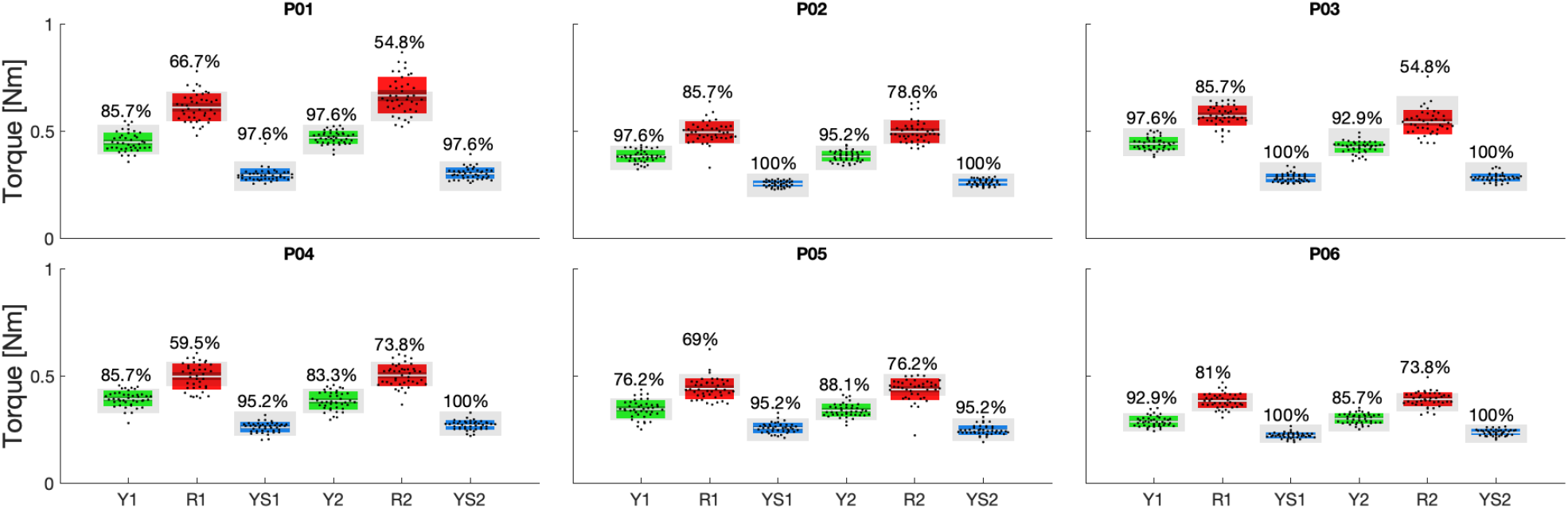
Participant-specific box plots reporting the distribution of torque response amplitude, with indications for the mean (white horizontal line), mean *±* one standard error of the mean (dark shaded area), and mean one *±* s.d. (light shaded area). The range indicating the cued torque response amplitude is provided in gray. Percent success rate for each task instruction is provided above the blox plots. 1 and 2 in the labels indicate 1^st^ or 2^nd^ task session.

Participants were generally successful when attempting to match torque target values, but the success rates were greater in yield than in the resist condition. Participants 1 and 3 had the lowest percentage success for Resist 2, though still significantly higher than Yield 2 at the individual participant level.

### B. Neural Correlates of LLRs

Fig. 6 shows the whole-brain group level imaging results. Generally, Yield and Resist activity localized in the contralateral primary, somatosensory, and premotor cortices. The threshold for statistical significance was identified as *t* = 12.84, calculated using SPM’s random-field theory correction. Clusters of highly significant activation for Yield localized in the ipsilateral visual cortex and the contralateral primary somatosensory cortex (Brodmann’s Area 2). Highly significant clusters of activation for Resist localized in the contralateral primary somatosensory cortex and the primary motor cortex (Brodmann’s Area 4). Additionally, the contrast for *β*_*R*_ *> β*_*Y S*_), indicating activity associated with the Resist condition while controlling for the Yield Slow condition (background activity only), resulted in clusters of highly significant activity in the contralateral primary motor cortex and premotor cortex (Brodmann’s Area 6).

**Fig. 6.**
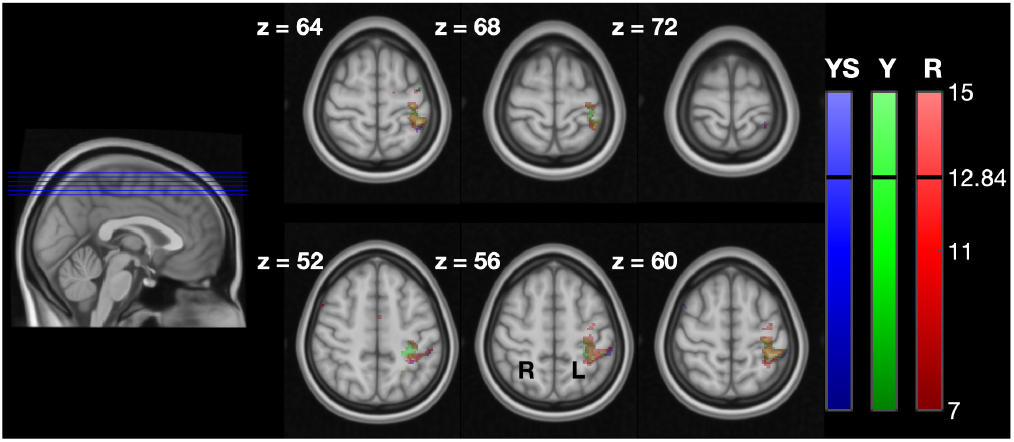
LLR-specific activation maps in the cortex for the different task instructions. The maps refer to contrasts *β*_*Y S*_ *>* 0, *β*_*Y*_ *>* 0, and *β*_*R*_ *>* 0 obtained for the whole-brain analysis. For each slice, the *z* coordinate is provided in mm.

Fig. 7 shows the brainstem-specific group level results. fMRI analysis revealed greater activation extent and intensity for the Resist condition compared to Yield, confirming the brainstem’s role in processing the task-dependent component of LLRs. Activation for Resist localized in the ipsilateral pons and bilaterally in the posterior medulla and midbrain. Yield activation was less widespread, localized in the ipsilateral pons and medulla, and contralateral midbrain. Given the small number of participants, the threshold for statistical significance was identified as *t* = 3, smaller than the threshold of *t* = 5.68 determined using small volume correction. A cluster of highly significant activation (*t >* 5.68) for Resist localized in the ipsilateral superior medulla at slice *z* = *−*32.

**Fig. 7.**
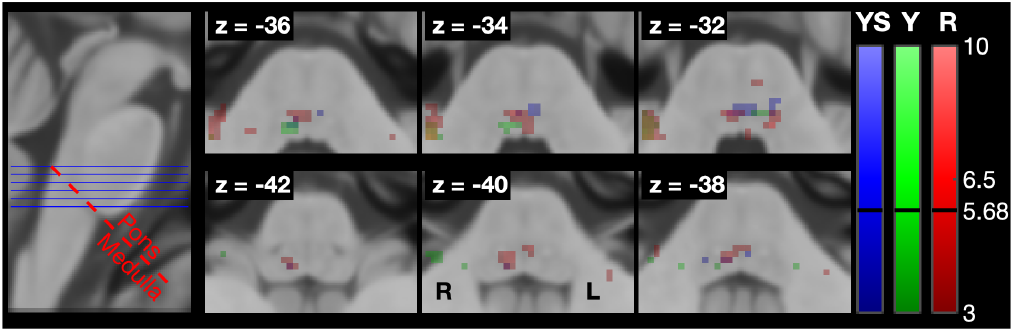
LLR-specific activation maps in the brainstem for the different task instructions, with overlaid the plane of demarcation used to refer to voxels as being in the pons or in the medulla. The maps refer to contrasts *β*_*Y S*_ *>* 0, *β*_*Y*_ *>* 0, and *β*_*R*_ *>* 0 obtained for the brainstem-specific analysis.

## IV. DISCUSSION

### A. Dual Motor StretchWrist and Task Paradigm

The DMSW, coupled with our task paradigm, offers a novel approach to elicit and condition LLRs. By providing participants with specific task instructions such as “Yield” or “Resist,” we can modulate LLRa. This capability is crucial for understanding the underlying neural mechanisms associated with LLRs and their adaptability based on different motor tasks.

While earlier investigations have employed the Yield-Resist paradigm, not all of them incorporated performance feedback for participants [13], [14]. Moreover, the majority of these studies exclusively utilized binary performance feedback [15], [16], [17]. Forgaard et al. extended this type of paradigm by utilizing angular displacement from a target as a form of performance feedback [18]. Our approach is similar as we condition performance feedback based on measured torque during the specified period, thereby enhancing the standardization of participants’ responses. This capacity to standardize participants’ torque responses throughout the conditioning process significantly contributes to the reliability and reproducibility of our experimental protocol.

However, our paradigm is novel in that we can perform the experiment in an MRI environment. In previous biofeedback-based paradigms, the perturbation was delivered as a torque perturbation, and feedback was provided based on the measured displacement. Due to constraints on MR-compatibility, the DMSW employs piezoelectric motors, which are a velocity source for the perturbations rather than a torque source. Our novel paradigm uses velocity perturbations, as allowed by our hardware, while using the measured torque to provide performance feedback. To our knowledge, it is the first time that these two modes have been combined to study the task-dependent modulation of LLRs.

Our preliminary imaging data have provided valuable insights into the neural correlates of LLRs under different task conditions. The observation of activity associated with “Resist” conditions in the contralateral primary motor cortex aligns with expectations, reflecting the engagement of cortical regions in response to the resistive motor task [19], [20], [21]. Moreover, the activation in the ipsilateral medulla and bilateral midbrain and pons aligns with previous studies’ findings on brainstem motor activity and suggests a robust involvement of the brainstem in processing the task-dependent components of LLRs [22], [23], [24]. This finding underscores the significance of considering both cortical and brainstem contributions in understanding motor responses.

### B. Limitations and Future Directions

Despite the promising results, there are limitations in our study. The small number of participants in both preliminary and pilot experiments, warrants caution in generalizing our findings. Future studies should aim to include a larger participant cohort to strengthen the external validity of our results.

Addressing the question of the suitability of our setup for stroke participants is critical. The observed limitations with individuals with higher body mass indexes and reduced capacity for movement and force production highlight the need for paradigm adjustments and considerations of different postures. Further research should explore modifications to accommodate stroke participants.

## V. CONCLUSIONS

We developed a new MRI-compatible robot, the DMSW, and validated a new experimental protocol based on the DMSW to evoke motor responses under different task instructions. Moreover, we found stretch response activity associated with the “resist” task instructions in the medulla. Future work will aim to establish the effect of different data processing strategies to improve the signal-to-noise ratio of fMRI of the brainstem, and to isolate the contributions of the brainstem to voluntary responses and LLRs.

## ACKNOWLEDGMENT

The authors would like to thank Cody Helm, Kristin Schmidt, and Manju Sivasankar for their assistance with participant data collections. The authors would also like to thank the Delaware Center for Biomedical and Brain Imaging.

## References

[1] J. G. McPherson, A. Chen, M. D. Ellis, J. Yao, C. J. Heckman, and J. P. Dewald, “Progressive recruitment of contralesional cortico-reticulospinal pathways drives motor impairment post stroke,” Journal of Physiology, vol. 596, pp. 1211–1225, apr 2018.

[2] C. YT, L. S, Z. P, and L. S, “A startling acoustic stimulation (SAS)-TMS approach to assess the reticulospinal system in healthy and stroke subjects,” Journal of the neurological sciences, vol. 399, pp. 82–88, apr 2019.

[3] C. F. Honeycutt, M. Kharouta, and E. J. Perreault, “Evidence for reticulospinal contributions to coordinated finger movements in humans,” 10.1152/jn.00866.2012, vol. 110, pp. 1476–1483, oct 2013.

[4] S. N. Baker, “The primate reticulospinal tract, hand function and functional recovery,” Journal of Physiology, vol. 589, pp. 5603–5612, ec 2011.

[5] S. Choudhury, A. Shobhana, R. Singh, D. Sen, S. S. Anand, S. Shubham, M. R. Baker, H. Kumar, and S. N. Baker, “The Relationship Between Enhanced Reticulospinal Outflow and Upper Limb Function in Chronic Stroke Patients,” Neurorehabilitation and Neural Repair, vol. 33, no. 5, pp. 375–383, 2019.

[6] J. Xu, A. M. Haith, and J. W. Krakauer, “Motor Control of the Hand Before and After Stroke,” Clinical Systems Neuroscience, pp. 271–289, jan 2015.

[7] N. A. Reddy, K. M. Zvolanek, S. Moia, C. Caballero-Gaudes, and M. G. Bright, “Denoising task-correlated head motion from motor-task fMRI data with multi-echo ICA,” Imaging Neuroscience, vol. 2, no. 2024, 2024.

[8] V. Singh, J. Pfeuffer, T. Zhao, and D. Ress, “Evaluation of spiral acquisition variants for functional imaging of human superior colliculus at 3T field strength,” Magnetic Resonance in Medicine, vol. 79, no. 4, pp. 1931–1940, 2018.

[9] I. L. Kurtzer, “Long-latency reflexes account for limb biomechanics through several supraspinal pathways.,” Frontiers in integrative neuroscience, vol. 8, p. 99, 2014.

[10] A. Zonnino, A. J. Farrens, D. Ress, and F. Sergi, “StretchfMRI: A novel technique to quantify the contribution of the reticular formation to long-latency responses via fMRI,” vol. 2019-June, pp. 1247–1253, 2019.

[11] G. N. Lewis, Æ. C. D. Mackinnon, and E. J. Perreault, “The effect of task instruction on the excitability of spinal and supraspinal reflex pathways projecting to the biceps muscle,” pp. 413–425, 2006.

[12] L. Kasper, S. Bollmann, A. O. Diaconescu, C. Hutton, J. Heinzle, S. Iglesias, T. U. Hauser, M. Sebold, Z. M. Manjaly, K. P. Pruessmann, and K. E. Stephan, “The PhysIO Toolbox for Modeling Physiological Noise in fMRI Data,” Journal of Neuroscience Methods, vol. 276, pp. 56–72, 2017.

[13] L. Spieser, S. Aubert, and M. Bonnard, “Involvement of SMAp in the intention-related long latency stretch reflex modulation: A TMS study,” Neuroscience, vol. 246, pp. 329–341, 2013.

[14] C. D. MacKinnon, M. C. Verrier, and W. G. Tatton, “Motor cortical potentials precede long-latency EMG activity evoked by imposed displacements of the human wrist,” Experimental Brain Research 2000 131:4, vol. 131, pp. 477–490, mar 2014.

[15] J. A. Pruszynski, I. Kurtzer, and S. H. Scott, “Rapid motor responses are appropriately tuned to the metrics of a visuospatial task,” Journal of Neurophysiology, vol. 100, no. 1, pp. 224–238, 2008.

[16] J. Weiler, P. L. Gribble, and J. A. Pruszynski, “Goal-dependent modulation of the long-latency stretch response at the shoulder, elbow, and wrist,” 10.1152/jn.00702.2015, vol. 114, pp. 3242–3254, ec 2015.

[17] L. Yang, J. A. Michaels, J. A. Pruszynski, and S. H. Scott, “Rapid motor responses quickly integrate visuospatial task constraints,” Experimental Brain Research, 2011.

[18] C. J. Forgaard, I. M. Franks, D. Maslovat, and R. Chua, “Influence of kinesthetic motor imagery and effector specificity on the long-latency stretch response,” 10.1152/jn.00159.2019, vol. 122, no. 5, pp. 2187–2200, 2019.

[19] R. A. Scheidt, J. L. Zimbelman, N. M. Salowitz, A. J. Suminski, K. M. Mosier, J. Houk, and L. Simo, “Remembering forward: Neural correlates of memory and prediction in human motor adaptation,” NeuroImage, vol. 59, no. 1, pp. 582–600, 2012.

[20] R. Shadmehr and J. W. Krakauer, “A computational neuroanatomy for motor control,” Experimental Brain Research, vol. 185, no. 3, pp. 359–381, 2008.

[21] A. J. Suminski, S. M. Rao, K. M. Mosier, and R. A. Scheidt, “Neural and electromyographic correlates of wrist posture control,” Journal of Neurophysiology, vol. 97, no. 2, pp. 1527–1545, 2007.

[22] A. Zonnino, A. J. Farrens, D. Ress, and F. Sergi, “Measurement of stretch-evoked brainstem function using fMRI,” Scientific Reports, vol. 11, no. 1, pp. 1–21, 2021.

[23] W. T. Chu, T. Mitchell, K. D. Foote, S. A. Coombes, and D. E. Vaillancourt, “Functional imaging of the brainstem during visually-guided motor control reveals visuomotor regions in the pons and midbrain,” NeuroImage, vol. 226, no. November 2020, 2021.

[24] V. J. Ravichandran, C. F. Honeycutt, J. Shemmell, and E. J. Perreault, “Instruction-dependent modulation of the long-latency stretch reflex is associated with indicators of startle,” Exp Brain Res, vol. 230, pp. 59–69, 2013.

